# Soluble Spike DNA vaccine provides long-term protective immunity against SAR-CoV-2 in mice and nonhuman primates

**DOI:** 10.1101/2020.10.09.334136

**Authors:** Yong Bok Seo, You Suk Suh, Ji In Ryu, Hwanhee Jang, Hanseul Oh, Bon-Sang Koo, Sang-Hwan Seo, Jung Joo Hong, Manki Song, Sung-Joo Kim, Young Chul Sung

**Author notes:** Correspondence and requests for materials should be addressed to Y.C.S.

## Abstract

The unprecedented and rapid spread of SARS-CoV-2 has motivated the need for a rapidly producible and scalable vaccine. Here, we developed a synthetic soluble SARS-CoV-2 spike (S) DNA-based vaccine candidate, GX-19. In mice, immunization with GX-19 elicited not only S-specific systemic and pulmonary antibody responses but also Th1-biased T cell responses in a dose-dependent manner. GX-19 vaccinated nonhuman primate seroconverted rapidly and exhibited detectable neutralizing antibody response as well as multifunctional CD4+ and CD8+ T cell responses. Notably, when the immunized nonhuman primates were challenged at 10 weeks after the last vaccination with GX-19, they did not develop fever and reduced viral loads in contrast to non-vaccinated primates as a control. These findings indicate that GX-19 vaccination provides durable protective immune response and also support further development of GX-19 as a vaccine candidate for SARS-CoV-2 in human clinical trials.

## Introduction

Severe acute respiratory syndrome coronavirus-2 (SARS-CoV-2) has emerged towards the end of 2019 as causative agent of pneumonia in the city of Wuhan in china (*1*). Since its emergence, the global situation is dynamically evolving, and on 30 January 2020 the World Health Organization declared Coronavirus Disease 2019 (COVID-19) as a public health emergency of international concern (PHEIC) and it was declared as a global pandemic on 11 March 2020. Disease symptoms range from mild flu-like to severe cases with life-threatening pneumonia (*2*). Unlike its predecessors of novel betacoronavirus such as SARS-CoV and MERS-CoV, SARS-CoV-2 transmits efficiently from person-to-person. Due to these high transmissibility and extensive community spread, SARS-CoV-2 has already caused nearly 20 million infection and over 700,000 death as of August 2020 (*3*). It is estimated that until ~60 to 70% of people develop herd immunity, it is unlikely to achieve sufficient control of SARS-CoV-2 to resume normal activities (*4*). Therefore, the benefit of the rapid development of vaccines against SARS-CoV-2 is very high and important to change the global epidemiology of this virus.

The four major structural proteins of SARS-CoV-2 are spike (S) protein, envelope (E) protein, membrane (M) protein, and nucleocapsid (N) protein, which is essential for the virus assembly and infection (*5*). The S protein is an attractive target for vaccine design because it plays crucial roles in receptor binding, fusion, and viral entry into the host cell during the infection process. Proteolytic cleavage of the S protein generates two subdomains, S1 and S2, and are responsible for host cell angiotensin-converting enzyme 2 (ACE2) receptor binding and host cell membrane fusion, respectively. While S2 domain is conserved across human coronaviruses, the S1 domain divergent across the coronaviruses (*6*). It has recently been reported that a vaccine strategy based on S antigen can prevent SARS-CoV-2 infection and disease in a mouse model (*7*). Moreover, studies in rhesus macaques have been shown that vaccine strategies based on the S antigen can prevent SARS-CoV-2 infection and disease in this relevant animal model (*8, 9*), indicating that the S antigen may be sufficient as a vaccine immunogen to elicit SARS-CoV-2 protective immunity.

The urgent need for vaccines prompted an international response, with the development of more than 170 candidate SARS-CoV-2 vaccine within the first 7 months of 2020 (*10*). In order to achieve an effective and rapid vaccine response against emerging viruses, a manufacturing and distribution platform that can shorten time to product availability as well as rapid vaccine design is important. While it typically takes more than several months to produce cell lines and clinical grade subunit proteins, nucleic acid vaccines can be completed in weeks (*11, 12*). In addition, DNA-based vaccines do not require a cold chain, making them a great alternative to the availability of important, life-saving vaccines in resource-poor areas of the world. There have been studies over the past decades to improve the efficacy of DNA vaccines, and as a result, many improvements in efficacy have been made (*13*). In addition to prophylactic DNA vaccines against viral infections such as MERS-CoV and Zika virus, a therapeutic DNA vaccine against cancers has been demonstrated in several clinical trials (*14–20*).

Here, we explored the potential of SARS-CoV-2 soluble S DNA vaccine candidate, designated GX-19. Vaccination with GX-19 elicits a concurrent humoral response and Th1-biased immune responses in both mice and non-human primate (NHP) models. Notably, the vaccine drives potent cellular and humoral response in NHPs, including neutralizing antibodies that provide potent protective efficacy against SARS-CoV-2 infection. The data demonstrate that the immunogenicity of this DNA vaccine supports the clinical development to advance the development of this DNA vaccine in response to the current global health crisis.

## Results

### Construction and immunogenicity of SARS-CoV-2 DNA vaccines

The representative sequence for the SARS-CoV-2 S protein was generated after analysis of the S protein genomic sequences, which were retrieved from the NCBI SARS-CoV-2 resources. The full-length or entire ectodomain of S gene was codon optimized for increased antigen expression in mammalian cells and N-terminal tissue plasminogen activator (tPA) signal sequence was added, respectively. The insert was then subcloned into the pGX27 vector (*21*) and the resulting plasmid was designated as pGX27-S and pGX27-S_ΔTM_ (Fig. 1A).

**Figure 1.**
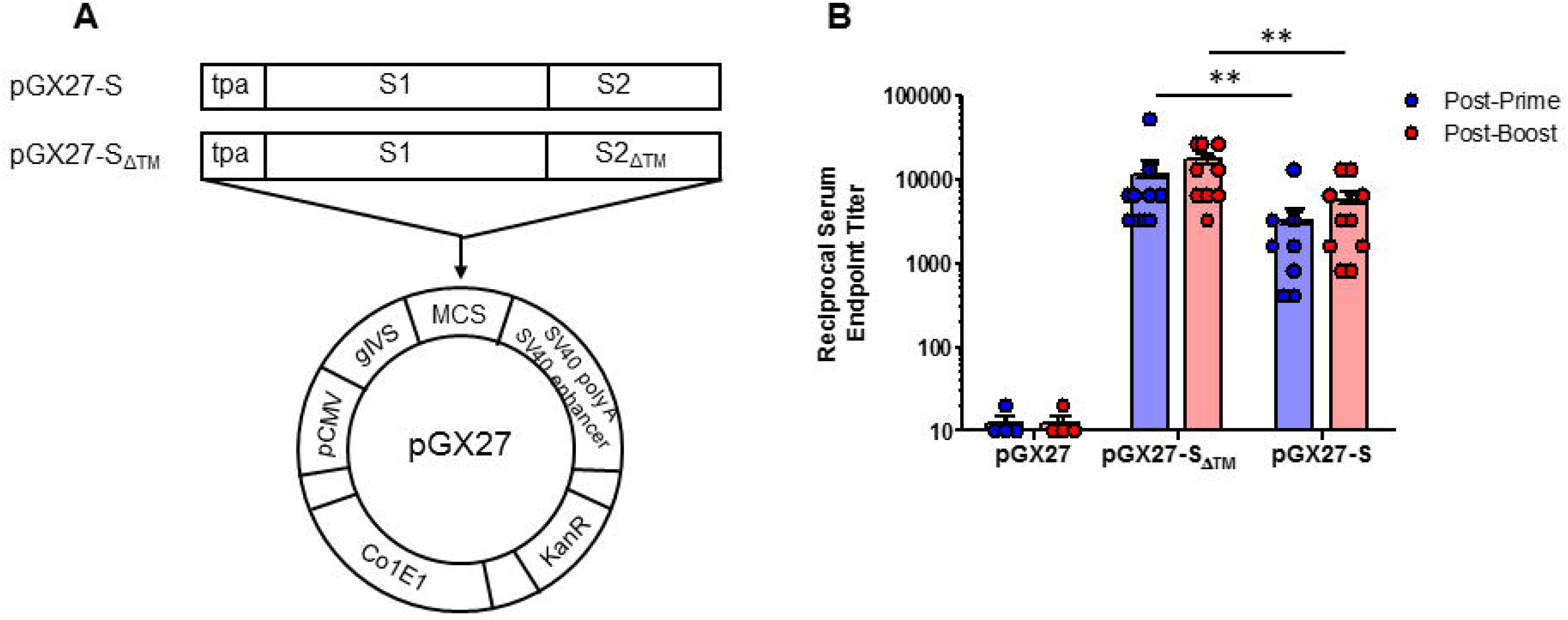
Diagram and immunogenicity of SARS-CoV-2 DNA vaccines. Schematic diagram of COVID-19 DNA vaccine expressing soluble SARS-CoV-2 S protein (S_ΔTM_) or full-length SARS-CoV-2 S protein (S) **(A)**. BALB/c mice (n=4-10/group) were immunized at week 0 and 2 with pGX27-S_ΔTM_, pGX27-S or pGX27 (empty control vector) as described in the methods. Sera were collected 2 weeks post-prime (blue) and 2 weeks post-boost (red) and evaluated for SARS-CoV-2 S-specific IgG antibodies **(B)**.

To assess the immunogenicity of two candidate DNA vaccines, we immunized six-week old female BALB/c mice intramuscularly (IM) twice at 2-week interval. As indicated Fig. 1B, immunized with the both of DNA vaccine candidate induced a robust S protein-specific antibody response compared to the control. Interestingly, there were higher antibody titers in the pGX27-S_ΔTM_ group than in the pGX27-S group at both post-prime and post-boost. These results were not consistent with the previous report that demonstrated higher binding antibody in full-length S DNA vaccinated macaques compared to S_ΔTM_ DNA vaccinated macaques (*8*).

### GX-19 induce strong humoral and cellular immune response in mice

pGX27-S_ΔTM_, named GX-19, was therefore selected as the vaccine candidate to advance to immunogenicity and efficacy studies. The immunization with GX-19 elicited significant serum IgG responses against the S protein in dose-dependent manner (Fig. 2A). Besides, GX-19 vaccination elicited S-specific IgG response in the bronchoalveolar lavage (BAL) fluid (Fig. 2B). Live virus neutralizing titers were also evaluated in BALB/C mice following the same GX-19 immunization regimen. Consistent to the S binding antibody response, potent neutralizing activity was elicited by GX-19 (Fig. 2C).

**Figure 2.**
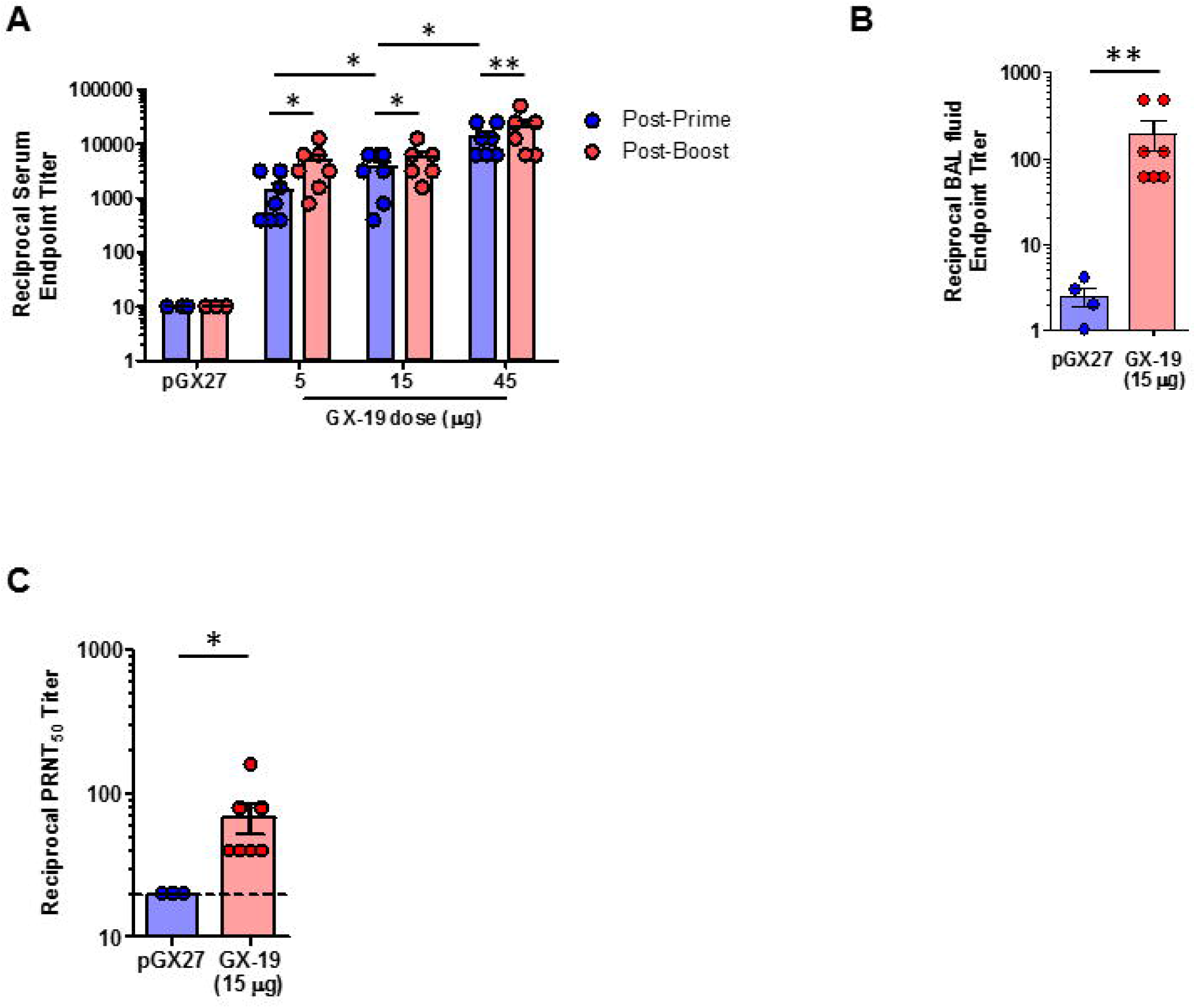
GX-19 elicit robust binding and neutralizing antibody responses in mice. BALB/c mice (n=4-7/group) were immunized at week 0 and 2 with indicated doses of GX-19 or pGX27 as described in the methods **(A-C)**. Sera were collected 2 weeks post-prime (blue) and 2 weeks post-boost (red) and assessed for SARS-CoV-2 S-specific IgG antibodies by ELISA **(A)**, and for post-boost sera, neutralizing antibodies against SARS-CoV-2 live-virus **(C)**. BAL were collected 2 weeks post-boost and assayed for SARS-CoV-2 S-specific IgG antibodies by ELISA **(B)**. Data representative of two independent experiments. *P* values determined by Mann-Whitney test.

Given that severe SARS-CoV is associated with Th2-biased immune response with inadequate Th1 response (*22–24*), we evaluated the balance of Th1 and Th2 response. Although BALB/c mice tend to develop a more Th2-biased response after vaccination (*25*). GX-19 induced Th1-biased immune response as indicated by higher IgG2a/b responses when compared to IgG1 regardless of vaccination doses (Fig. 3A and 3B). We also directly measured cytokine patterns in vaccine-induced T cells by cytometric bead array. GX-19-induced T cells secreted large amount of IFN-γ, TNF-α, or IL-2 but did not secrete IL-4 or IL-5 (Fig. 3D). At 2 weeks after the boost vaccination, the spleens were harvested and S-specific T cell response were measured by IFN-γ ELISPOT. Mice immunized with 5-, 15-, 45-μg of GX-19 exhibited dose-dependent splenic T cell response with mean IFN-γ spots per 10^6^ cells of 542, 872, and 1932, respectively (Fig. 3C). To gain further insight into the responses of GX-19-induced T cell responses, we also measured multiple cytokines by intracellular cytokine staining (ICS) as described in Methods. GX-19 vaccination exhibited significantly higher percentage of S-specific CD4^+^ T cells or CD8^+^ T cells secreting IFN-γ, TNF-α, and IL-2 (Fig. 3E; Supplementary Fig. 1).

**Figure 3.**
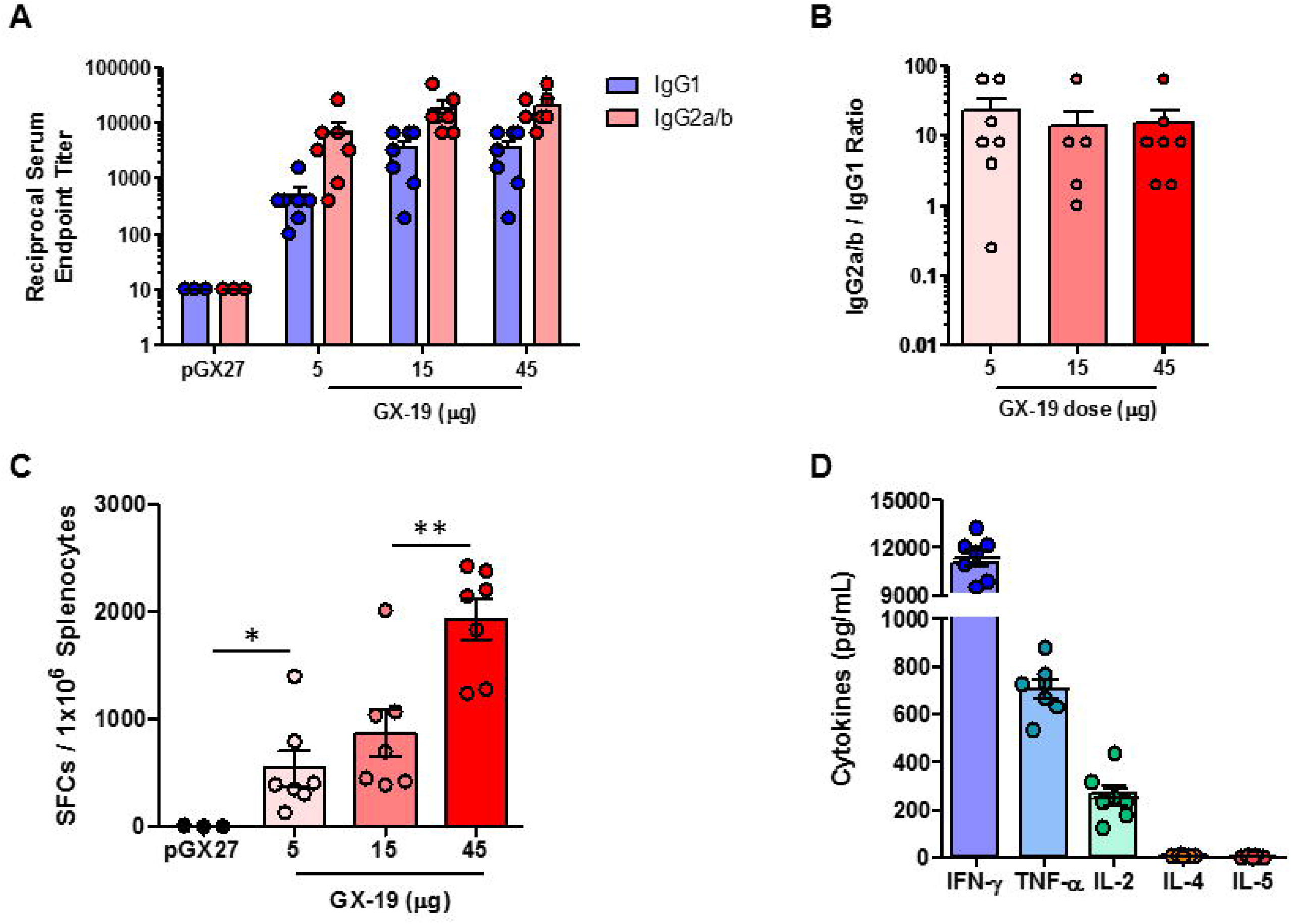
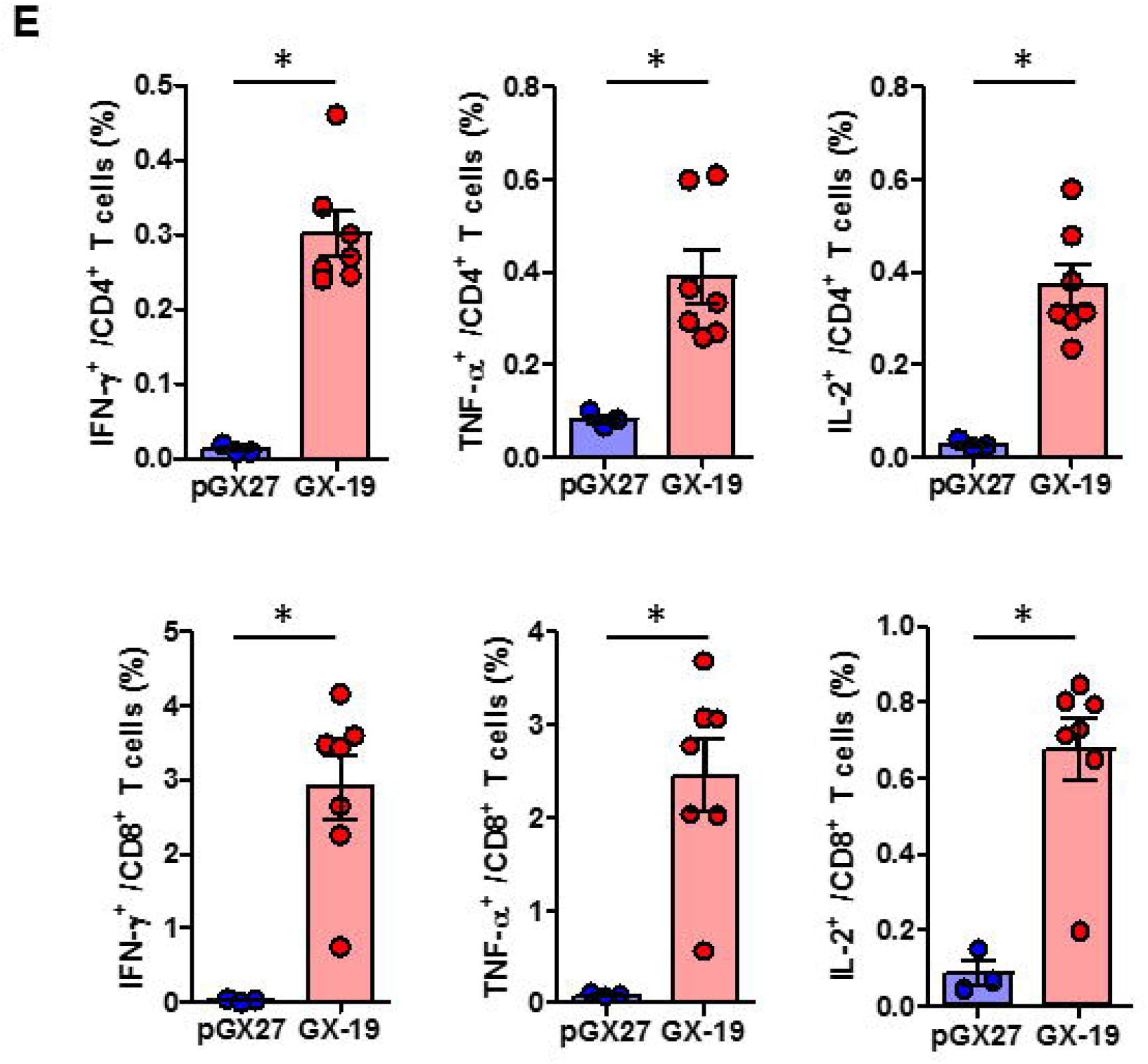
Immunization with GX-19 elicit Th1-biased T cell responses in mice. BALB/c mice (n=3-7/group) were immunized at week 0 and 2 with indicated doses of GX-19 or pGX27 (empty control vector) as described in the methods **(A-C)**. Sera were collected 2 weeks post-boost and assessed for SARS-CoV-2 S-specific IgG1 and IgG2a/b. Endpoint titers **(A)**, and endpoint tier ratios of IgG2a/b to IgG1 **(B)** were calculated. 2 weeks post-boost mouse splenocytes were isolated and re-stimulated with peptide pools spanning the SARS-CoV-2 S protein *ex vivo*. Indicated cytokines in the supernatants of culture were quantified using Th1/Th2 cytometric bead array kit **(D)**. T cell responses were measured by IFN-γ ELISPOT in splenocytes stimulated with peptide pools spanning the SARS-CoV-2 S protein **(C)**. Cells were stained for intracellular production of IFN-γ, TNF-α, and IL-2. Shown are the frequency of S-specific CD4^+^ or CD8^+^ T cells after subtraction of background (DMSO vehicle) **(E)**. Data representative of two independent experiments. *P* values determined by Mann-Whitney test.

### GX-19 elicit robust humoral and cellular immune response in NHPs

To investigate whether GX-19 was capable of inducing immune responses in a NHP model, three macaques were vaccinated with electroporation (EP)-enhanced delivery three times about 3-week intervals with GX-19, as described in Methods. Blood was collected at week-0, −5.5, and −8 to monitor vaccine-induced immune responses. The binding ELISA results showed that all three macaques immunized with GX-19 seroconverted after a single immunization with anti-S IgG titers tending to increase by boosting vaccination (Fig. 4A). In addition, sera collected at week-0, −5.5 and −8 were further analyzed for neutralization of wild-type SARS-CoV-2 (Korea CDC) by the 50% plaque reduction neutralization test (PRNT_50_). Three macaques immunized with GX-19 displayed elevated neutralization titers with mean PRNT_50_ titers of 1: 285 (at week 5.5) and 1: 996 (at week 8), respectively (Fig. 4B).

**Figure 4.**
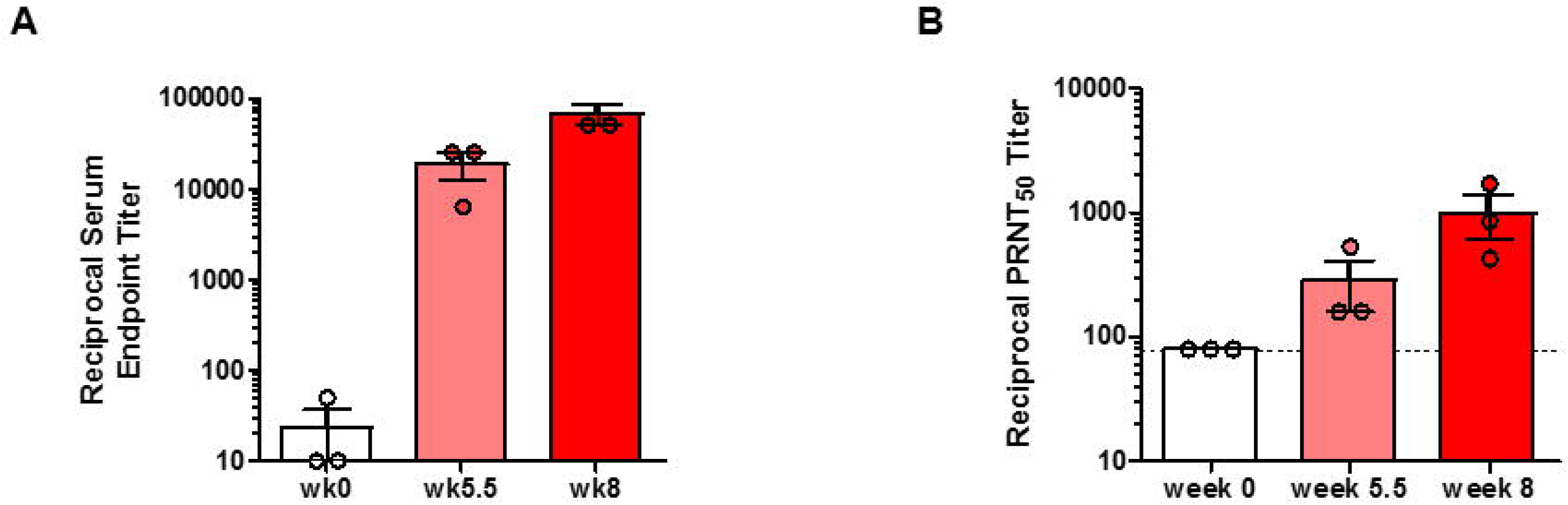
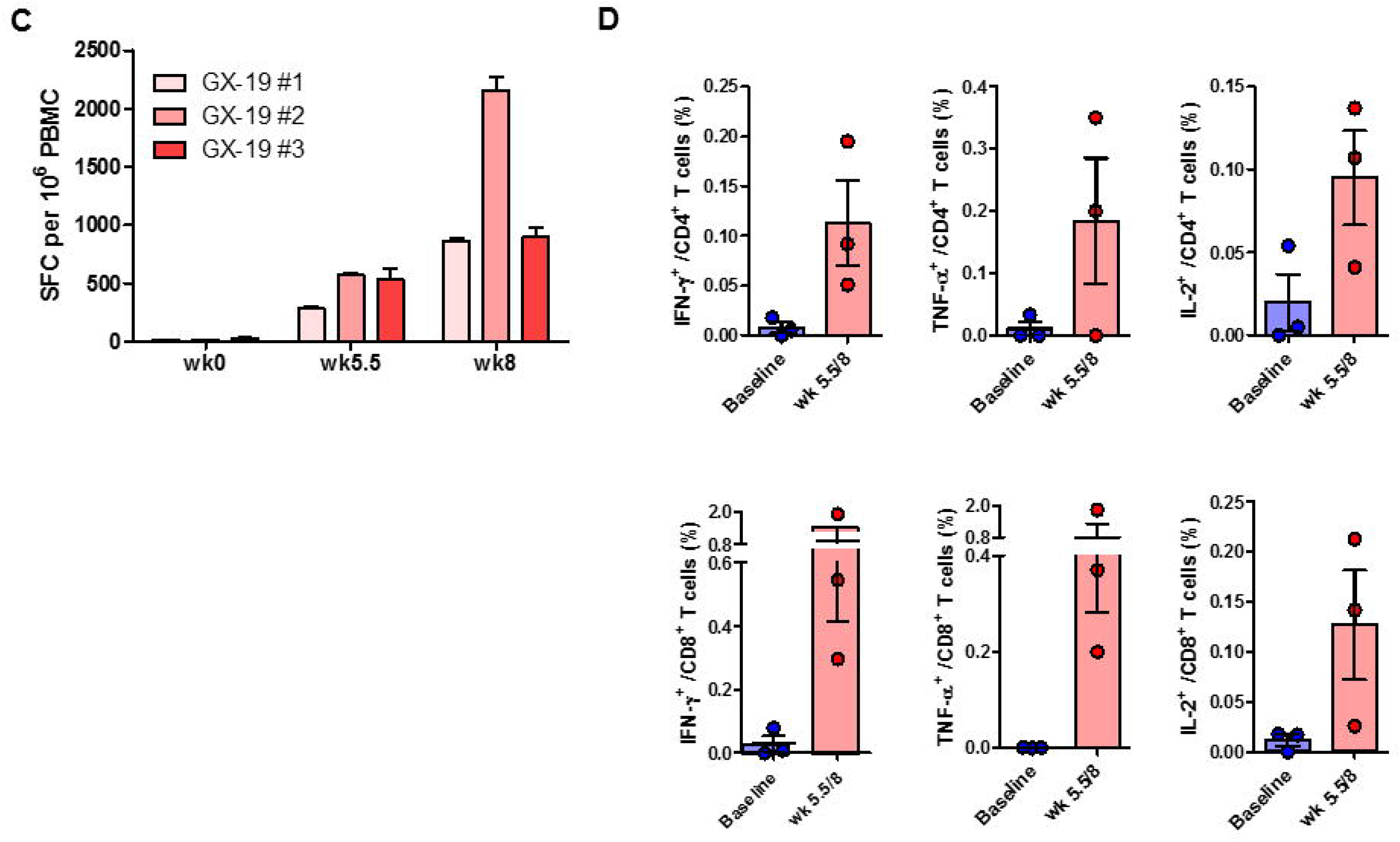
Antibody and T-cell responses after GX-19 vaccination in macaques. Macaques (n=3) were immunized with 3 mg of GX-19 as described in the methods. Serum and PBMCs were collected before (wk 0), during (wk 4, and 5.5) and after (wk 8) vaccination and were assessed for SARS-CoV-2 S-specific IgG antibodies by ELISA **(A)** and neutralizing antibodies against SARS-CoV-2 live-virus **(B)**. Data represent mean SEM of individual macaques, and dashed line indicate the assay limits of detection. The number of SARS-CoV-2 S-specific IFN-γ secreting cells in PBMCs was determined by IFN-γ ELISPOT assay after stimulation with peptide pools spanning the SARS-CoV-2 S protein. Shown are spot-forming cells (SFC) per 10^6^ PBMCS in triplicate wells **(C)**. The frequency of S-specific CD4^+^ or CD8^+^ T cells producing IFN-γ, TNF-α, or IL-2 was determined by intracellular cytokine staining assays stimulated with SARS-CoV-2 S peptide pools. Shown are the frequency of S-specific CD4^+^ or CD8^+^ T cells after subtraction of background (DMSO vehicle) **(D)**.

To determine the impact of the GX-19 on cellular immune response, ELISPOT analysis was used to measure T cell responses in blood of the vaccinated macaques. Three macaques developed T cell responses after single immunization. In addition, all animals exhibited such an elevated responses after boost vaccination indicating vaccine-induced cellular immune responses became progressively stronger in macaques during GX-19 vaccination (Fig. 4C). To gain further insight into the responses of GX-19-induced T cell responses, we also measured multiple cytokines by ICS as described in Methods. GX-19 vaccination exhibited meaningful induction S-specific CD4^+^ T cells or CD8^+^ T cells producing IFN-γ, TNF-α and, to a lesser extent, IL-2 (Fig. 4D; Supplementary Fig. 2).

### GX-19 protect NHPs from wild-type SARS-CoV-2 infection

Induction of long-term immunological memory for T cell and B cell responses is important for effective vaccine development. Unlike other vaccine studies in which NHPs confirmed protective efficacy against SARS-CoV-2 infection within 4 weeks after the last vaccination (*8, 9, 26, 27*), we evaluated the protective efficacy approximately 10 weeks after the last vaccination. To assess the protective efficacy of GX-19, macaques were challenged by multiple routes with a total dose of 2.6 × 10^7^ 50% tissue-culture infectious doses (TCID_50_), on 10 weeks after the last vaccination. This challenge route and dose were based on a model development study in which we challenged macaques that had no previous exposure to the virus (*28*). GX-19 vaccinated macaques showed no increase in body temperature after viral infection, and showed rapid recovery in lymphocyte reduction compared to unvaccinated macaques (Supplementary Fig. 3A and 3B). High levels of viral load were observed in the unvaccinated macaques (Fig. 5A and 5B) with a median peak of 7.54 (range 6.66 - 8.01) log_10_ viral copies/ml in nasal swab and a median peak of 6.18 (range 6.01 - 6.81) log_10_ viral copies/ml in throat swab (Fig. 5C and 5D). Although there is no statistical significance due to insufficient number of animals, lower levels of viral load were observed in GX-19 vaccinated macaques, including 1.58 and 1.57 log_10_ reductions of median peak viral load in nasal swab and throat swab, respectively (Fig 5A - D). Since the peak viral load does not reflect the presence of total virus over time, the virus load then calculated based on the area under curve (AUC). GX-19 vaccinated macaques had a viral AUC of 6.02 ± 0.23 log_10_ in nasal swab and 4.99 ± 0.45 log_10_ in throat swab, respectively, which were 1.46 and 1.45 log_10_ decreases of the viral AUC compared to unvaccinated macaques respectively (Fig. 5E and 5F). To confirm for infectious viruses, TCID_50_ assay was performed for nasopharyngeal swab and oropharyngeal swab samples on Vero cells. The infectious viral load showed the similar pattern as viral RNA load. Although there is no statistical significance due to insufficient number of animal, lower levels of infectious viral load were observed in GX-19 vaccinated macaques (Supplementary Fig. 3C and 3D).

**Figure 5.**
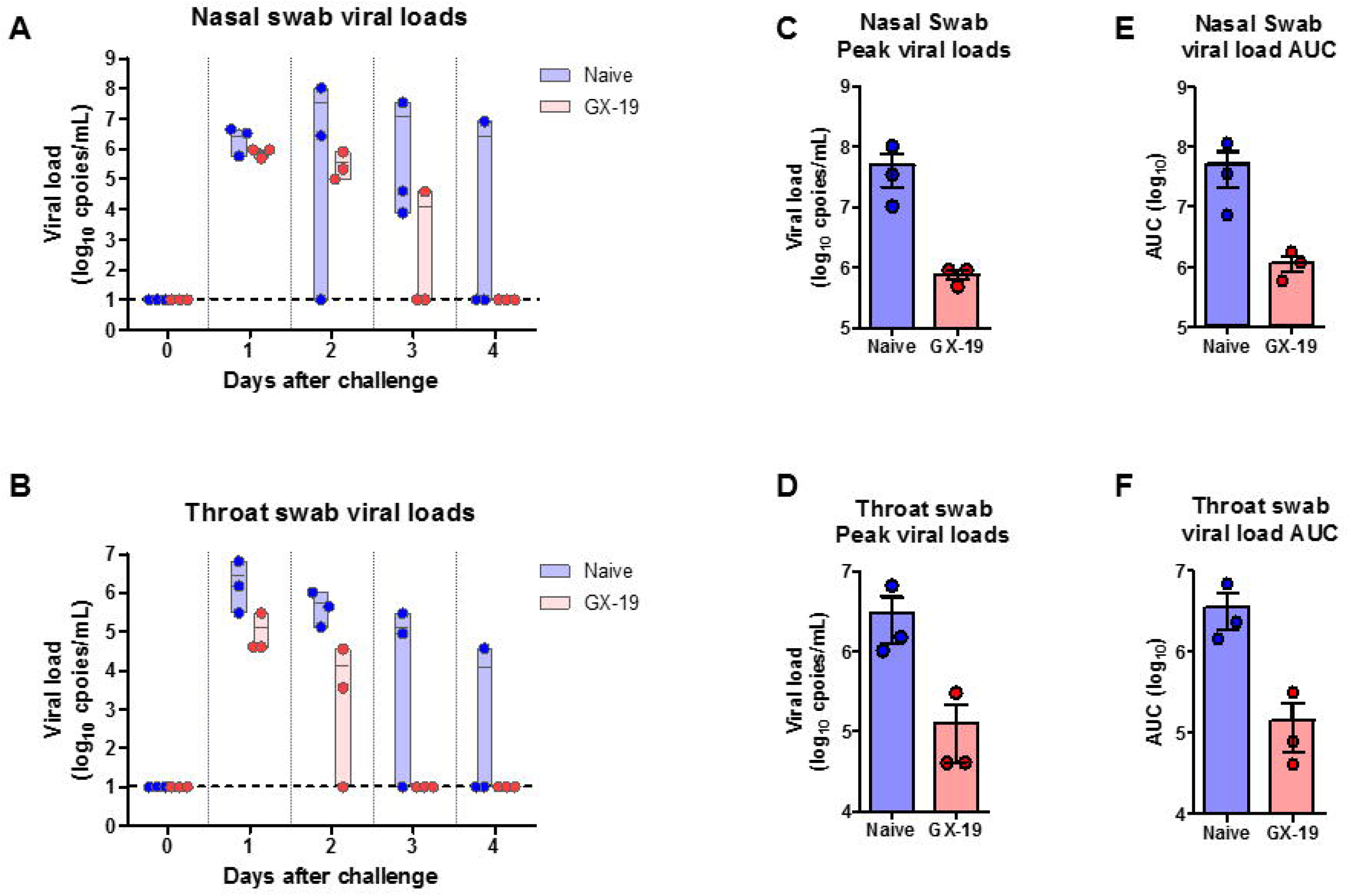
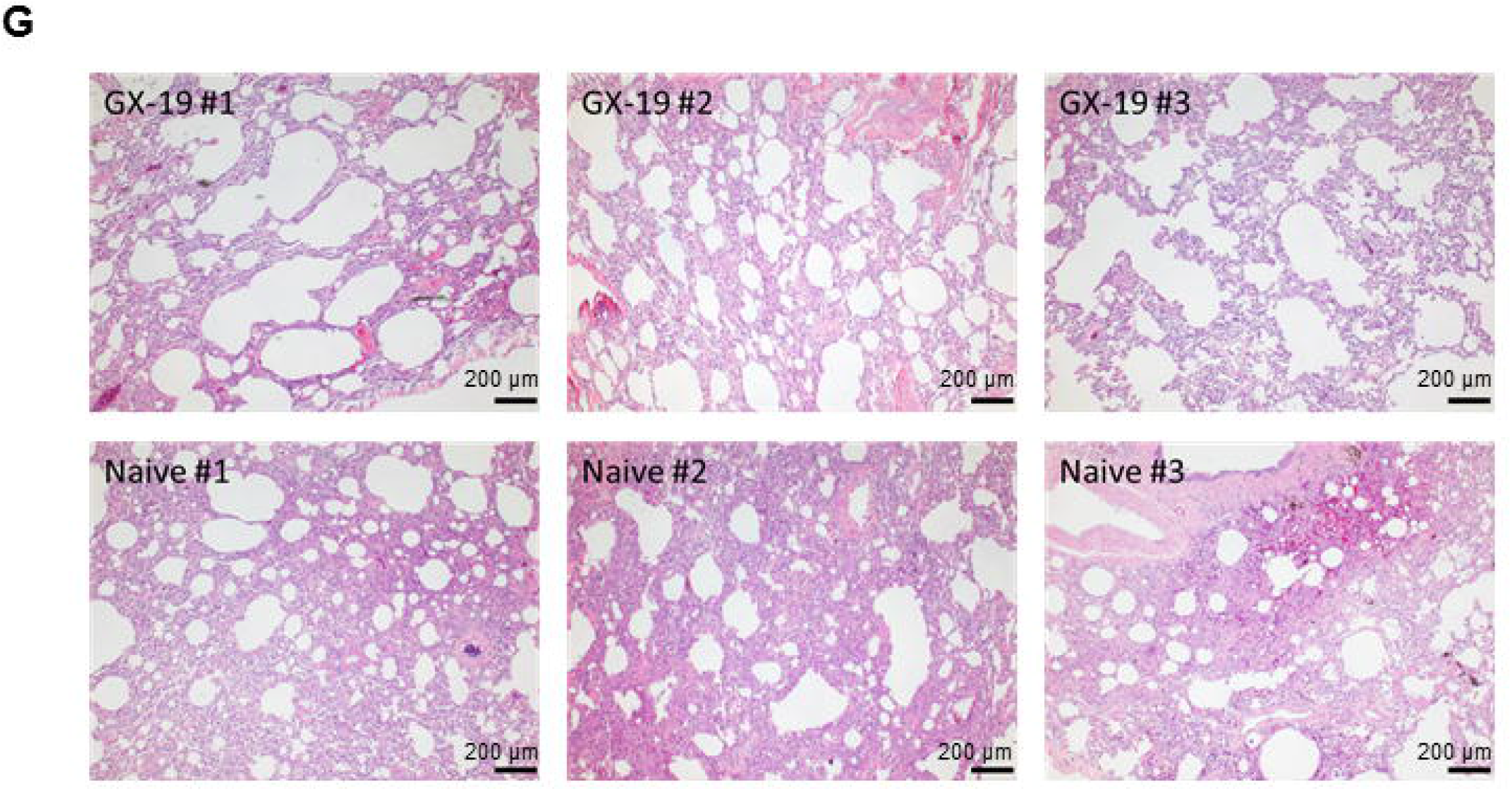
Protective efficacy of GX-19 against SARS-CoV-2 challenge. Non-vaccinated (n=3) and GX-19 vaccinated macaques (n=3) were challenged by intratracheal, oral, conjunctival, intranasal, and intravenous administration of 2.7 × 10^7^ TCID_50_ SARS-CoV-2. Viral load were assessed in nasal swab **(A)** and throat swab **(B)** at multiple time-points following challenge. Summary of peak viral loads and viral load AUC in nasal swab **(C, E)** and throat swab **(D, F)** following challenge. Dashed line indicate the assay limit of detection. Histopathological changes in the lungs of SARS-CoV-2 challenged macaques **(G)**. The lung tissue sections were stained with hematoxylin and eosin (H&E).

At 4 days post virus inoculation, all animals were euthanized, and tissues were collected. Consistent with previous reports (*28*), SARS-CoV-2 infection caused moderate-to-severe inflammation, as evidenced by small airways and the adjacent alveolar interstitia in non-vaccinated macaques. In vaccinated macaques, the viral challenge caused mild histopathologic changes compared to those in control macaques (Fig. 5G).

## Discussion

In this study, we demonstrated that GX-19 (pGX27-S_ΔTM_) exhibited higher S-specific antibody response than pGX27-S. In addition, GX-19 can elicit SARS-CoV-2 S-specific Th1-biased T cell response in mice and NHPs. Vaccination of GX-19 can confer effective protection against SARS-CoV-2 challenge at 10 weeks following the last vaccination.

In a recent study, the low immunogenicity and protective efficacy of S_ΔTM_ was reported in the evaluation of the protective efficacy of DNA vaccine expressing various forms of SARS-CoV-2 S protein (*8*). In contrast to full-length S DNA here, we showed that GX-19 not only induces excellent antibody responses in mice and NHPs, but also effective protection against SARS-CoV-2 virus challenge in NHPs. The different results can be explained by the difference in the cellular localization of antigen and the difference in the strength of the induced immune responses by vaccination. In a study on the effects of cellular and humoral immune responses according to cellular localization of antigens after DNA immunization, it was found that the DNA encoding secreted OVA produces a much higher immune response than the cytoplasmic or membrane-bound form (*29*). In the previous study (*8*), the vaccines were administered without EP method which significantly enhanced the *in vivo* delivery efficacy of DNA vaccine by 100 - 1,000 folds (*30*), and as a result, they induced a weak antibody responses or T cell responses. On the other hand, electroporation-enhanced GX-19 induced robust antibody and T cell responses. In addition, the strength of the immune response increases depending on the strength of the expression vector (*31*), and pGX27 vector has about three-time higher expression strength than commercial vector (unpublished data). Here, we believe that the introduction of a high-expression vector (*21*) into GX-19 along with an effective EP delivery system resulted in the efficient protective effect against SARS-CoV-2 infection through the induction of a strong immune responses. However, further studies will be needed to compare pGX27-S and pGX27-S_ΔTM_ in NHPs under our conditions to confirm this finding.

In this study, we observed that GX-19 induced concurrent antibody, CD4^+^ T, and CD8^+^ T cell response in both mice and NHPs models. Indeed, successful DNA vaccination effectively induces complete complementation of the immune responses, including humoral and cellular responses (CD8^+^ and Th1 cellular responses) similar to those achieved by live attenuated viruses (*14, 15, 32, 33*). This can be explained by the nature of DNA vaccine, presumably because both class-I antigen-processing pathways (i.e., intracellular processing of viral proteins into peptides and subsequent loading onto MHC class-I molecules) and class-II antigen-processing pathways (i.e., specifically engineered in the S signal sequence that cause an increased export of viral surface antigens) are possible. Among T cell responses, the balanced Th1/Th2 responses are important because vaccine-associated enhanced respiratory disease (VAERD) is associated with Th2-biased immune response. Indeed, immunopathologic complications characterized by Th2-biased immune responses have been reported in animal model of the SARS-CoV or MERS-CoV challenge (*22, 34–38*), and similar phenomena have been reported in clinics vaccinated with whole-inactivated virus vaccines against RSV and measles virus (*39, 40*). In addition, the importance of T cell responses has been highlighted by recent study of asymptomatic and mild SARS-CoV-2 convalescent (*41*). These results collectively suggest that vaccines capable of generating balanced antibody responses and T cell responses may be important in providing protection against SARS-CoV-2 diseases. Here, we show that GX-19 induces Th1-biased responses, suggesting DNA vaccination can avoid Th2-biased immune response associated with VARED. In fact, there were subtle pathologic changes in the SARS-CoV-2 infected GX-19 vaccine group. These results were also demonstrated in the other respiratory infection DNA vaccine such as SARS-CoV and MERS-CoV (*42*), MERS-CoV (*32*). This suggests that DNA vaccine platform can be a good alternative in vaccine development for emerging infections where the balanced T cell response as well as antibody response is important.

It is desirable that the SARS-CoV-2 vaccine can prevent infection or disease and induce long-term immunity. Virus-specific T cell responses play an important role in antiviral and disease control. Immune-modulatory cytokines (e.g., IFN-γ, TNF-α, and IL-2) released from virus-specific CD4^+^ T and CD8^+^ T cells play a key role in several antiviral responses and act in synergy with type I IFNs to inhibit viral replication (*43, 44*). Patients with impaired IFN-γ activity were reported to have 5-fold increased susceptibility to SARS (*45*). Clinical cases of asymptomatic virus infection indicate that virus-specific T cells can control disease even in the absence of neutralizing antibodies (*41, 46*). Here, we showed that GX-19 induce potent antigen-specific CD4^+^ and CD8^+^ T cell activation and robust release of immune-modulatory cytokines in mice and NHPs, indicating that GX-19 can effectively control the disease of SARS-CoV-2. Next, clinical cases in which antibody responses rapidly decrease and disappear after SARS-CoV-2 infection indicate the importance of vaccines that can induce long-term immunological memory (*47–49*). GX-19 induced potent CD4^+^ and CD8^+^ T cells in both animal models and it may confer long-lasting immunity against coronaviruses as indicated in SARS survivors, where CD8^+^ T cell immunity persisted up to 11 years (*43, 50*). Although the results of the long-term immune response were not covered here, we observed an effective protection against viral infection about 10 weeks after the last vaccination, indicating that the GX-19-induced memory immune response is also effective against long-term infection. In summary, these results explain the promising immunogenicity of the GX-19 and, in particular, support the clinical potential of the GX-19 by showing the protective effect against infection from NHPs.

## Methods

### DNA vaccine construction

SARS-CoV-2 DNA construct encodes a consensus SARS-CoV-2 spike (S) protein sequences developed by comparing the sequences of current reported sequences. The vaccine insert was codon optimized and commercially synthesized (Thermo Fisher Scientific). Synthetic genes were subcloned into high-expression vector, pGX27 (*21*).

### Mouse immunizations

Female BALB/c mice aged 6-8 weeks (Koatech) were immunized with GX-19 vaccine or pGX27 in a total volume of 50 μl of PBS into the tibialis anterior muscle with in vivo electroporation with OrbiJector^®^ (SL VAXiGEN Inc.) at weeks 0, 2. On day 14 and 28 blood or BAL (bronchoalveolar lavage) fluid was collected and mice were sacrificed 14 days after final immunization.

### NHP immunizations and challenge

Cynomolgus macaques that weighed between 3.4 and 4.4 kg and had a mean age of 5.6 years were immunized 3 mg GX-19 vaccine at week 0, week 3 and week 5.5. GX-19 vaccine was given intramuscularly, followed by electroporation using OrbiJector^®^ (SL VAXiGEN Inc.). Blood was collected immediately before the first immunization (week 0), and every 1-3 weeks thereafter through week 8 and sera and PBMCs were isolated to evaluate the humoral or cellular immune response, respectively. At 10 weeks after the last vaccination, immunized macaques were moved to the animal biosecurity level 3 (ABL-3) laboratory in the Korea National Primate Research Centre (KNPRC) at the Korea Research Institute of Bioscience and Biotechnology (KRIBB). All animals were challenged with total 2.6 × 10^7^ 50% tissue culture infectious doses/mL [TCID_50_]/mL SARS-CoV-2 virus, obtained from the National Culture Collection for Pathogens (accession number 43326), via combined routes (intratracheal, oral, conjunctival, intranasal, and intravenous route) as previously described (*28*). After viral challenge, macaques were anesthetized with a ketamine sodium (10 mg/kg) and tiletamine/zolazepam (5 mg/kg) at 0, 1, 2, 3, and 4 days post-infection (dpi) and conducted the following procedure: checking body temperature, weight, and respiration rate and collecting nasopharyngeal, oropharyngeal swab samples in universal transport medium. Swab samples were centrifuged at 1600 x g for 10 min and filtered with 0.2 μm pore size syringe filters for further virus quantification. Immunization procedures were approved by the Institutional Animal Care and Use Committee (IACUC permit number ORIENT-IACUC-20044), and challenging procedures were approved by KRIBB IACUC (permit number KRIBB-AEC-20178).

### Antigen binding ELISA

Serum and BAL fluid was collected at each time point was evaluated for binding titers. Ninety-six well immunosorbent plates (NUNC) were coated with 1 μg/mL recombinant SARS-CoV-2 S1+S2 ECD protein (Sino Biological 40589-V08B1), S1 protein (Sino Biological 40591-V08H) in PBS overnight at 4°C. Plates were washed 3 times with 0.05% PBST (Tween 20 in PBS) and blocked with 5% skim milk in 0.05% PBST (SM) for 2-3 hours at room temperature. Sera or BAL fluid were serially diluted in 5% SM, added to the wells and incubated for 2 hours at 37°C. Following incubation, plates were washed 5 times with 0.05% PBST and then incubated with horseradish peroxidase (HRP)-conjugated anti-mouse IgG (Jackson ImmunoResearch Laboratories 115-035-003), IgG1 (Jackson ImmunoResearch Laboratories 115-035-205), or IgG2a (Jackson ImmunoResearch Laboratories 115-035-206) or IgG2b (Jackson ImmunoResearch Laboratories 115-035-207) for the mouse sera/BAL or anti-monkey IgG (Bethyl Laborabories A140-102P) for the NHP sera for 1 hour at 37°C. After final wash plates were developed using TMB solution (Surmodics TMBW-0100-01) and the reaction stopped with 2N H_2_SO_4_. The plates were read at 450 nm by SpectraMax Plus384 (Molecular Devices).

### Live virus neutralization assay

1.5 × 10^4^ Vero cells were seeded on 96-well plate (NUNC 167008) and incubate at 37°C and 5% CO_2_ for 16 hours. 25 μl of diluted sera were mixed with SARS-CoV-2 (NCP43326) (300 plaque-forming unit per 25 μl) and incubated the mixture at 37°C and 5% CO_2_ for 30 minutes. The cell culture media of the Vero cells were removed and the Vero cells were washed using 200 μl of serum-free DMEM (Invitrogen 11995065). The virus-sera mixture was treated into the Vero cells and incubated at 37°C and 5% CO_2_ for 4 hours. After incubation, the treated mixture was removed and the cells were washed using 100 μl of phosphate buffered saline (PBS) (Gibco 10010-023). The Vero cells were fixed using 300 μl of 10% formalin solution (Sigma F8775) by incubating at 4°C for overnight. After washing, the Vero cells were permeabilized by adding 100 μl of ice-cold 100% methanol (Sigma D7). After 10 minutes incubation at room-temperature, the methanol was removed and the Vero cells were washed using 100 μl PBS, blocked using 100 μl of blocking buffer (0.5% normal goat serum (Abcam Ab7481) + 0.1% Tween 20 (GenDEPOT T9100-100) + 1% (w/v) Bovine serum albumin (Sigma A3803-100G) in PBS) and incubated at room temperature for 30 minutes incubation at room temperature. After removing the blocking buffer, 3,000-fold diluted 100 μl of anti-SARS-CoV-2 NP rabbit mAb (Sino Biological 40143-R001) was added on the Vero cells and incubated at 37°C for 1 hour. After removing the Ab solution on the Vero cells, the Vero cells were washed using 200 μl of PBS containing 0.1% Tween 20. 2,000-fold diluted goat anti-rabbit IgG-HRP (Bio-Rad 170-6515) solution was treated on the Vero cells and incubated at 37°C for 1 hour. After removing the goat anti-rabbit IgG-HRP solution, the Vero cells were washed using 200 μl of PNS containing 0.1% Tween 20. 30 μl of TrueBlue solution was added on the Vero cells and incubate at room temperature for 30 minutes. After removing the TrueBlue solution, the cells were air dried until completely dry. The numbers of focus of each well were read using CTL reader (Cellular Technology Ltd.) and the neutralizing Ab titers were calculated using Microsoft Excel and SoftMax (Version 5.4.1.).

### IFN-γ ELISPOT

For mouse samples, the Mouse IFN-γ ELISPOT set (BD 551083) was used as directed by the manufacturer. ELISPOT plates were coated with purified anti-mouse IFN-γ capture antibody and incubated overnight at 4°C. Plates were washed and blocked for 2 hours with RPMI + 10% FBS (R10 media) Five hundred thousand splenocytes were added to each well and stimulated for 24 hours at 37°C in 5% CO_2_ with R10 media (negative control), concanavalin A (positive control), or specific peptide antigens (2 μg/ml). Peptide pools consisted of 15-mer peptides overlapping by 11 amino acids and spanned the entire SARS-CoV-2 S protein (GenScript). After stimulation, the plates were washed and spots were developed according to the manufacturer’s instructions. For NHP samples, the Monkey IFN-γ ELISpot^PLUS^ kit (MABTECH 3421M-APT-10) was used as directed by the manufacturer. Two hundred thousand PBMCs were stimulated with peptide pools, and plates were washed and spots were developed according to the manufacturer’s instructions. Plates were scanned and counted on AID ELISPOT reader classic. Spot-forming unit (SFU) per million cells was calculated by subtracting the negative control wells.

### Intracellular cytokine staining

For mouse samples, splenocytes were stimulated in R10 media with specific peptide pools or medium alone (DMSO control) for 12 hours. After stimulation, cells were washed with PBS for subsequent immunostaining. Antibodies for staining cells were CD8 FITC (Biolegend 100706), IL-2 PE (Biolegend 503808), CD4 PE-Cy7 (Biolegend 100528), IFN-γ APC (Biolegend 505810), TNF-α (Biolegend 506328), CD3 BV605 (Biolgend 100351), and Live/dead IR (Invitrogen L10119). For NHP samples, cryopreserved and thawed PBMCs were resuspended in R10 media, and rested overnight at 37°C, 5% CO_2_, and subsequently PBMCs were stimulated in R10 media with specific peptide pools or medium alone (DMSO control) in the presence of 1 μg/ml of α-CD28 (BD bioscience 555725) and α-CD49d (BD bioscience 555501) for 12 hours. After stimulation, cells were washed with PBS for subsequent immunostaining and polychromatic flowy cytometric analysis. Antibodies for staining cells were CD3 PE (BD bioscience 552127), CD4 PerCP-Cy5.5 (Biolegend 317428), CD8a PE-Cy7 (Biolegend 301012), IFN-γ APC (Biolegend 506510), TNF-α BV421 (Biolegend 502932), IL-2 BV605 (Biolegend 500332), and Live/dead Near-IR (Invitrogen L10119). Fluorescence-activated cell sorting analysis was accomplished by Fortessa flow cytometer (BD bioscience), and the data were analyzed using FlowJo software. Background cytokine expression in the DMSO-controls was subtracted from that measured in the S peptide pools.

### Cytokine profile analysis by cytometric bead array (CBA)

Five hundred thousand splenocytes were plated and stimulated in R10 media with peptide pools (15-mers with 11-mer overlaps) corresponding to the SARS-CoV-2 S proteins (2 μg/ml) or the medium only as negative control in 96-well plates. Culture supernatants were harvested 48 hours after the stimulation and cytokines were quantitated by the BDTM CBA Mouse Th1/Th2 Cytokine kit (BD Biosciences) according to manufacturer’s instructions.

### Virus identification and quantification

Filtered swab samples were inoculated into Vero cells and incubated for 3 days at 37°C, for virus isolation to calculate the values of TCID50/mL using the Reed and Muench method. The viral RNA genome was extracted from the supernatant using QIAamp Viral RNA Mini Kit (Qiagen) and stored at −80°C in the ABL-3 facility until use. RT-qPCR was performed with a primer set targeting partial regions of the ORF1b gene in the SARS-CoV-2 virus using the QIAGEN OneStep RT-PCR kit (Qiagen) as previously reported (*51*). For all RT-qPCR analyses, SARS-CoV-2 RNA standard and negative samples were run in parallel for determination of virus copy number.

### Histological evaluation

Six lobes of the lung samples (three lobes in the right and left lung, namely upper, middle and lower lobe) of infected macaques were fixed in 4% paraformaldehyde for a minimum of 7 days, embedded in paraffin, and 4- to 5-μm sections were stained with hematoxylin and eosin.

### Statistical analysis

Analysis of virologic and immunologic data was performed using GraphPad Prism 5 (GraphPad Software). Comparison of data between groups was performed using two-sided Mann-Whitney tests. *P* values <0.05 were considered significant.

---

**Supplementary Figure 1. Flow cytometry panel to quantify SARS-CoV-2 S-specific mouse CD4^+^ or CD8^+^ T cells**.

Mouse splenocytes were stimulated with specific peptide pools and then analyzed with multicolor flow cytometry to simultaneously detect SARS-CoV-2 S-specific expression of IFN-γ, TNF-α, and IL-2. Gating strategy to identify CD4^+^ or CD8^+^ T cells (A). The representative plots show the frequencies of IFN-γ, TNF-α, IL-2 producing CD4^+^ or CD8^+^ T cells, respectively (B).

**Supplementary Figure 2. Flow cytometry panel to quantify SARS-CoV-2 S-specific NHP CD4^+^ or CD8^+^ T cells**.

Cryopreserved PBMCs of GX-19 vaccinated macaques were stimulated with specific peptide pools and then analyzed with multicolor flow cytometry to simultaneously detect SARS-CoV-2 S-specific expression of IFN-γ, TNF-α, and IL-2. Gating strategy to identify CD4^+^ or CD8^+^ T cells (A). The representative plots show the frequencies of IFN-γ, TNF-α, IL-2 producing CD4^+^ or CD8^+^ T cells, respectively (B).

**Supplementary Figure 3. Clinical signs and TCID_50_/mL in macaques challenged with SARS-CoV-2 after vaccination with GX-19**.

Non-vaccinated and vaccinated macaques (n=3/group) were challenged with 2.7 × 10^7^ TCID_50_ SARS-CoV-2. After viral challenge, macaques were anesthetized for checking body temperature (A), and blood lymphocyte count (B) at 0, 1, 2, 3, and 4 days post-infection (dpi). TCID_50_/mL was measured in nasopharyngeal (C), and oropharyngeal (D) at multiple time-points following challenge.

## Acknowledgment

The pathogen resources (NCCP43326) for this study were provided by the National Culture Collection for Pathogens.

