## Supplementary figures and images for "Soluble Spike DNA vaccine provides long-term protective immunity against SAR-CoV-2 in mice and nonhuman primates"

### Supplementary Figure 1

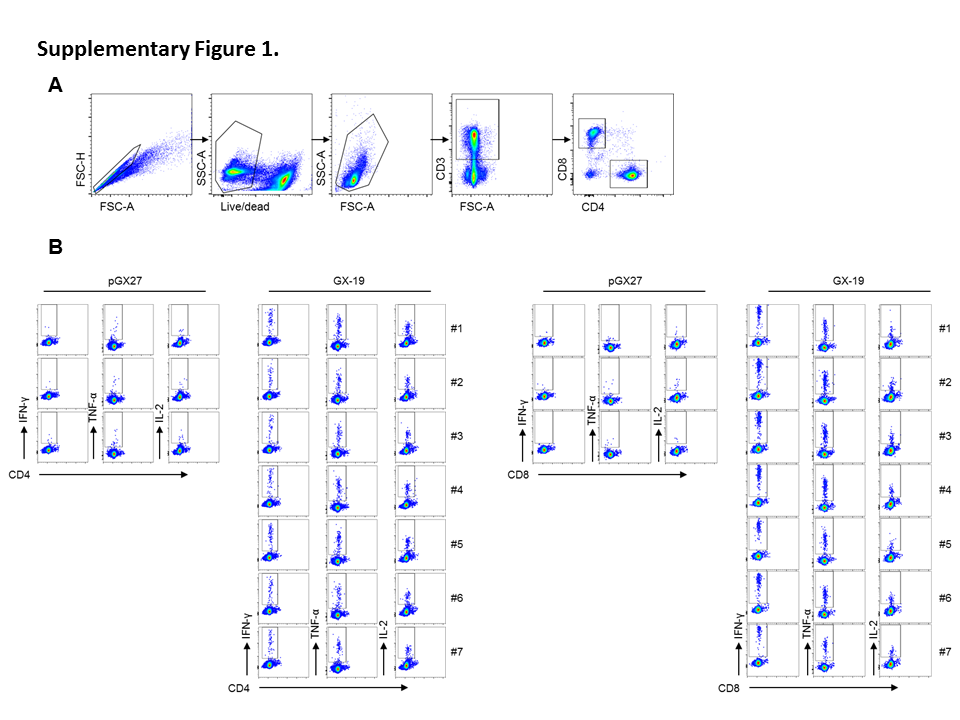

### Supplementary Figure 2

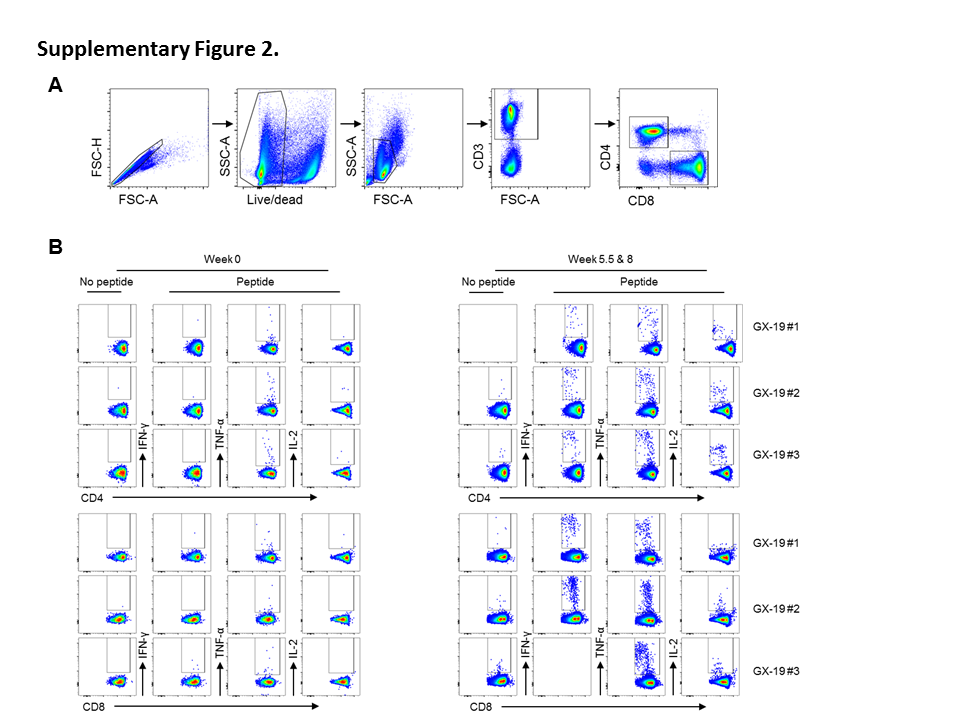

### Supplementary Figure 3

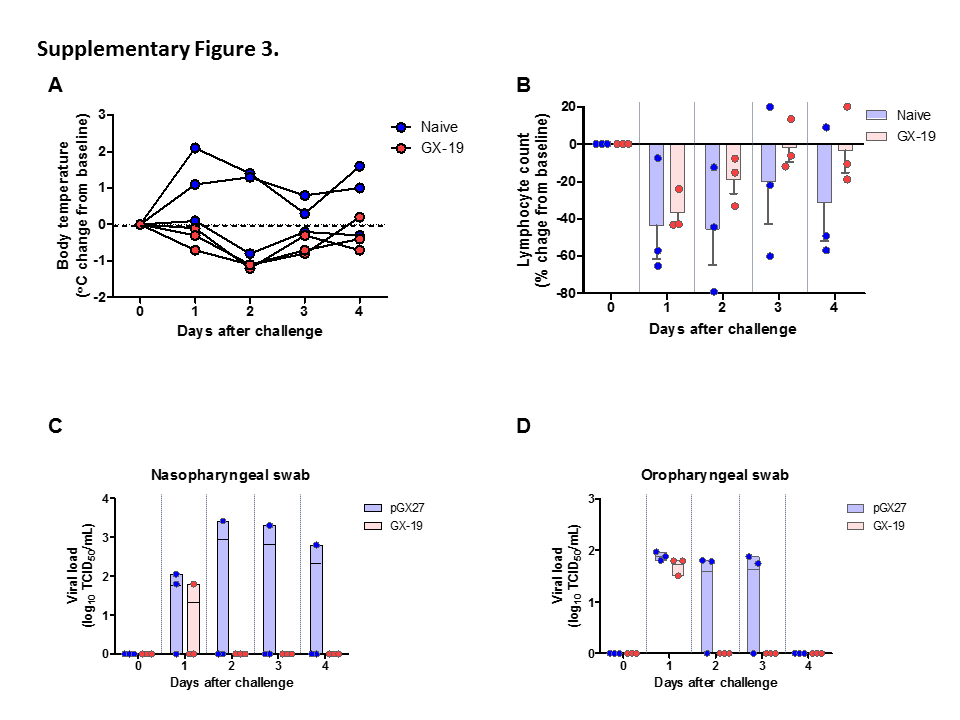
